# Alternative Conformations of lncRNAs Identified Through Structural Deconvolution of SHAPE- and DMS-MaP Datasets

**DOI:** 10.1101/2024.08.06.606861

**Authors:** Lucy Fallon, Alisha N Jones

## Abstract

The biological function of many classes of RNAs depend on their structures, which can exist as structural ensembles, rather than a single minimum free energy fold. In the past decade, long noncoding RNAs (lncRNAs) have emerged as functional transcripts in gene regulation that behave through their primary sequences and the structures they adopt. Chemical probing experiments, like selective 2’-hydroxyl acylation analyzed by primer extension and mutational profiling (SHAPE-MaP), and dimethyl sulfate-MaP (DMS-MaP), facilitate the characterization of RNA secondary structure both inside and outside the cell. But chemical probing experiments yield an *average* reactivity profile, representative of all the structures a particular RNA transcript adopts at the time of chemical probing, weighted by their relative populations. Chemical probing experiments often struggle to identify coexisting conformations a lncRNA might sample. Computational methods (DRACO, DREEM, DANCE-MaP) have been developed to identify alternate conformations of RNAs by deconvoluting chemical probing data. In this work, we investigate the propensity for lncRNAs to sample multiple structured states, and find each of the studied lncRNAs possess coexisting folds. We discuss the implications of lncRNAs harboring multiple structures and how it may contribute to their multifunctionality in regulating biological processes.

## Introduction

It has become increasingly evident that long noncoding RNAs (lncRNAs), which are a class of RNA transcripts longer than 500 nucleotides (nt) and lack protein-coding potential, have key regulatory roles in a wide repertoire of biological processes, including gene expression, chromatin remodeling, and epigenetics^1–4^. It is through their sequences and structural folds that lncRNAs interact with RNA binding proteins (RBPs) and other nucleic acids, acting as decoys, scaffolds, guides, and sponges to regulate biological function^5^. For example, a 2016 study reported the secondary structure of the lncRNA XIST (X-inactive specific transcript) both *in vitro* and in cells, demonstrating how the structural organization of XIST is modulated upon binding by RBPs^6^. Another study focusing on the interactions between the lncRNA nuclear enriched abundant transcript 1 (NEAT1) and several RBPs, revealed how binding of the RBP NONO (non-POU domain containing octamer) induces higher-order oligomerization, leading to phase separation and the formation of paraspeckles^7^.

An interesting, but understudied aspect in lncRNA biology is the ability of a lncRNA to behave as a multifunctional transcript; that is, a single transcript can regulate numerous biological functions. The functional intergenic repeating RNA element (FIRRE) lncRNA, for example, has regulatory roles in both adipogenesis and hematopoiesis^8–10^. Considering that many lncRNAs function through interactions with RBPs, and that many lncRNAs are bound by scores of RBPs, this supports a rising hypothesis that lncRNA structural heterogeneity may enable lncRNAs to bind multiple proteins such that they can behave as multifunctional transcripts^11,12^. Indeed, emerging studies have suggested that lncRNAs are conformationally dynamic, with their structures being best described as an *ensemble* of structures, instead of a single static folded state^6,13–15^. Each structured state is linked to a biological process. For example, the lncRNA COOLAIR samples different conformations depending on if it is exposed to warm or cold temperatures, and the different conformations modulate transcription activity and flowering^16^. In addition to temperature, other factors that can influence the structural dynamics of RNAs include pH, interactions with RNA binding proteins (RBPs), small molecules, ions, DNA, other RNAs, and even post-transcriptional modifications^17–19^.

To date, several thousands of biologically relevant lncRNAs have been identified in humans alone; the propensity by which RNAs sample multiple conformations, however, is still largely unknown^20^. In this study, we apply a computational method for the deconvolution of RNA alternative conformations (DRACO) to publicly available *in vitro* and in-cell selective 2’-hydroxyl acylation analyzed by primer extension and mutational profiling (SHAPE-MaP) and dimethyl sulfate-MaP (DMS-MaP) chemical probing datasets to identify coexisting conformations for five structurally-characterized lncRNAs: XIST, SLNCR1 (SRA-like noncoding RNA 1), GAS5 (growth-arrest specific lncRNA 5), MALAT1 (metastasis associated lung adenocarcinoma transcript 1), and MEG3 (maternally expressed gene 3)^6,21–24^. We show that some regions of these lncRNAs, which have been previously reported to adopt a single static structure, instead sample multiple coexisting conformations. These conformationally dynamic regions are (i) often sites of RBP binding, and (ii) occur locally, rather than globally within a lncRNA. We discuss how the conformational dynamics of lncRNAs contributes to their multifunctionality, and how interactions between lncRNAs and other biomolecules may help modulate lncRNA structure. Our results are further evidence for the growing recognition that lncRNAs are conformationally dynamic, and suggest that the different conformations lncRNAs adopt are associated with different biological functions. Future work will further explore the relationship between specific lncRNA conformations and their biological function.

## Methods

### Benchmarking and applying DRACO to SHAPE-MaP and DMS-MaPseq experiments

Since DRACO was designed for DMS-MaP experiments – not SHAPE-MaP – we first demonstrate that DRACO is capable of recovering known conformational states of the human immunodeficiency virus (HIV) Rev Response Element (RRE) from SHAPE-Map experiments^25^. The HIV-1 RRE was *in vitro* transcribed with bacteriophage T7 RNA polymerase, prepared in-house. In short, 0.64 μM DNA template was supplemented with 20 mM MgCl_2_, 8 mM each rATP, rCTP, rGTP, rUTP, 5% polyethylene glycol 8000, 1X transcription buffer (5 mM Tris pH 8, 5 mM spermidine, 10 mM DTT), and 0.6 mg of T7 RNA polymerase. Succeeding a 1 hour induction at 37°C, HIV-1 RRE was purified by high performance liquid chromatography (HPLC) at a gradient of 500 to 0 mM NaClO_4_. Isolated RNA was then dialyzed in a gradient, 1 M to 0 M, NaCl solution, and dehydrated by lyopholization.

*In vitro* SHAPE experiments were performed on the purified HIV-1 RRE using NMIA. 5 pmol of RNA was combined with 6 μL H_2_O, and refolded in 3.3X buffer (333 mM HEPES (pH 8.0), 333 mM NaCl and 33 mM MgCl_2_.The chemical probing was conducted at 40°C, with an NMIA concentration of 5 mM and 9 μL of refolded RNA. Modifications were mapped onto the RRE sequence by reverse transcription using SuperScript™ II Reverse Transcriptase and following the workflow as has been described in a previous study^26^.

To identify optimal parameters for deconvoluting coexisting conformations of the RRE (and other lncRNA transcripts) from SHAPE-MaP data, we used the RNAFramework suite to map the reads, count the mutations, and generate files for deconvolution with DRACO ^25,27^. The forward and reverse reads were first merged using NGmerge^28^. The merged reads were mapped to the HIV RRE reference transcript using the *rf-map* module of RNAFrameworks (flags *-b2 –bs –cqo –cq5 20 –ctn –cmn 0 –mp “––very-sensitive-local”*), and the *rf-count* module was then used (flags *-r –m –ds 200 –es –nd –ni –me 0.1 –dc 3 –mm*) to count the mutations and generate the mutation map files for deconvolution with DRACO. The mutation map files were then combined, and were deconvoluted using DRACO (flags *––shape ––absWinLen 100*)^25^. The outputted .json file was then processed using *rf-json2rc* with default parameters, and the returned .rc file was normalized with the *rf-norm* module (flags *-sm 4 –nm 2*). Finally, the *rf-jackknife* module was used to determine optimal slope and Y-intercept values for folding, which were then used in the *rf-fold* module to fold the deconvoluted structures.

DRACO could recover the previously reported 5– and 4 stem-loop (SL) structures of the HIV-1 RRE, validating that DRACO can successfully deconvolute SHAPE-MaP data (**Supplementary** Figure 1). Using the same approach, we identified coexisting conformations in five lncRNAs (XIST, SLNCR1, GAS5, MALAT1, and MEG3) by deconvoluting publicly available *in vitro* and in cell SHAPE– / DMS-MaP experiments. Their lengths, respective gene IDs, transcript IDs, and the source of the chemical probing data used for deconvolution are provided in **Supplementary Table 1**. The same input parameters for the HIV RRE were used for the five lncRNAs, only altering the flags that distinguish between SHAPE– and DMS-MaP datasets.

RNA structures were visualized and curated for illustration using RNACanvas^29,30^.

## Results and Discussion

### Deconvoluting coexisting conformations in five lncRNAs

DRACO is one of several computational algorithms that can detect structurally dynamic regions in an RNA. Following mutational profiling chemical probing experiments, DRACO identifies clusters of sequenced reads based on their co-mutation patterns^16,25,31–33^. Each cluster represents a distinct conformation sampled by the RNA; the relative number of reads in each cluster can be used as a proxy for the relative population of structures in the cell.

### The lncRNA XIST

The XIST lncRNA is one of the most well-studied lncRNA transcripts to date, and has been extensively reviewed from both a functional and structural perspective^34–40^. The human homologue of XIST is ∼19 kb long (∼18 kb in mouse); this transcript acts as the master regulator for X-chromosome inactivation (XCI) in placental mammals^37,39,41^. It is composed of six repetitive domains: the A-, F-, B-, C-, D– and E-repeats. The mouse XIST A-repeats have been structurally characterized in over 15 studies, based on computational and experimental approaches, and interestingly, are not in agreement^6,14,38,42–50^. Broadly, the reported structures of the XIST A-repeats can be categorized into modular and non-modular folds^50^. The disparity among different reports is an indication that the A-repeats may sample multiple structured states^17,50^. Inspection of the structural models proposed for the XIST A-repeat domain reveals paired nucleotides with high chemical probing reactivities, as well as single-stranded nucleotides with low reactivities, which is an indicator of conformational heterogeneity^17^. To confirm this, we used DRACO to investigate the conformational dynamics of the A-repeats.

The only published dataset with a sufficient number of reads for mouse XIST structural deconvolution comes from the Weeks research group^6^. Deconvolution of their 1-methyl-7-nitroisatoic anhydride (1M7) chemical probing data revealed i) high Shannon entropies for several regions across the length of the transcript, which indicates a high degree of structural heterogeneity and ii) one large region within the mouse XIST A-repeats that is conformationally dynamic (**Figure 1A, B**). This region is predicted to sample two different conformations. The conformationally dynamic region alternates between a more branched, modular structure – or a more linear, extended structure, represented at a 0.25:0.75 ratio, respectively. The structures predicted by DRACO contain elements that resemble each of the structures that have been formerly proposed. A more in depth analysis, using higher resolution approaches (similar to unpublished work on the structural organization of the XIST A-repeats by Jones, et al) is necessary to obtain more detailed information on the A-repeat structural ensemble^17^. Nevertheless, these results ascertain that the XIST A-repeats are conformationally dynamic.

**Figure 1.**
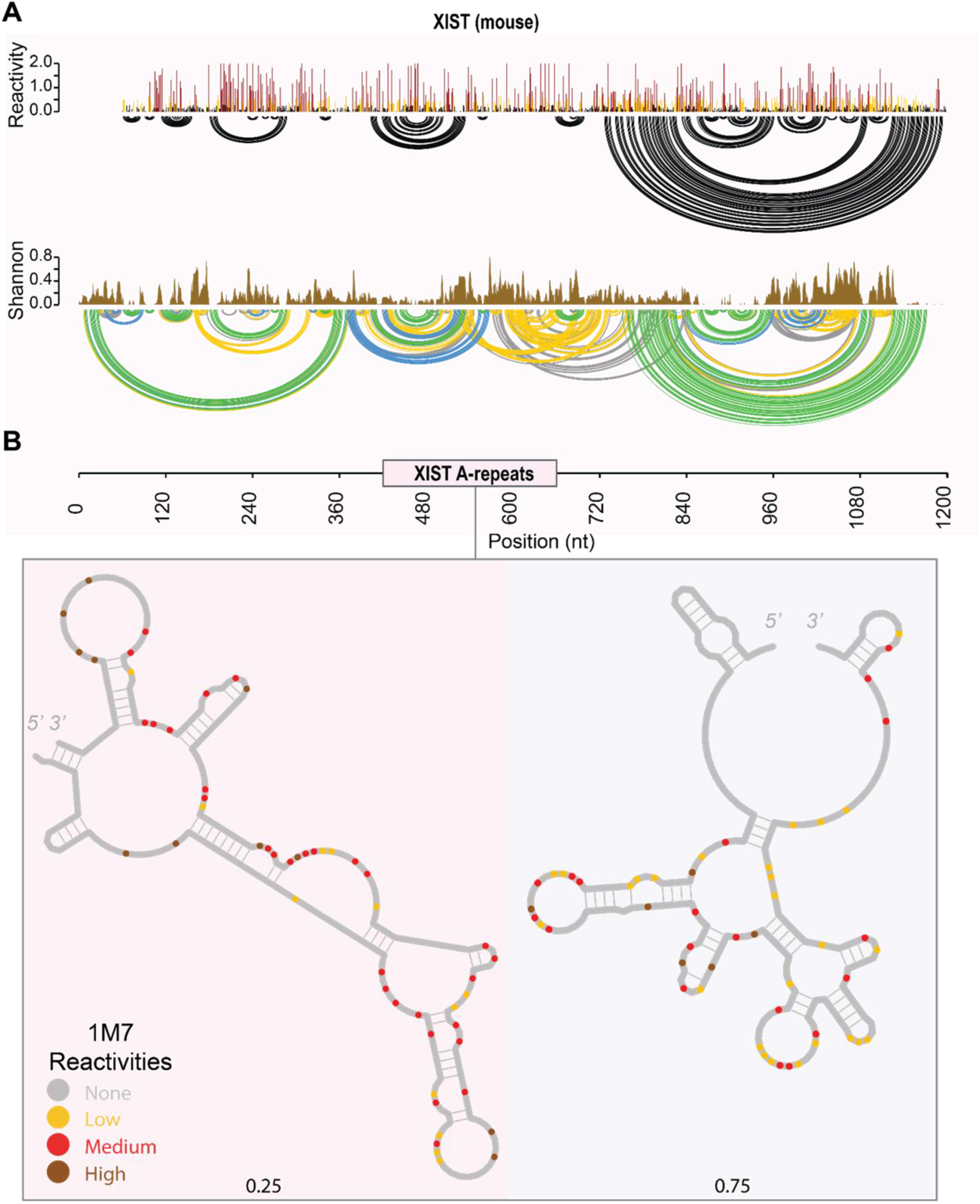
Deconvolution of the mouse Xist lncRNA using the Weeks chemical probing dataset^6^. **A.** Reactivities, arc plot, and Shannon entropy of the mouse Xist A-repeats. **B.** One region within the A-repeats of mouse Xist was predicted by DRACO to sample coexisting conformations. The deconvoluted structures are represented with their respective stoichiometries and deconvoluted reactivities.

The mouse A-repeats demonstrate conformational heterogeneity; these dynamics occur locally, as opposed to global rearrangements of structure. This may be due to the finite window size utilized by DRACO during deconvolution^25^. Interestingly, the A-repeats are bound by nearly 15 different RBPs, each with distinct RNA sequence and structural binding preferences^6,47,50–63^. The structural dynamics associated with the A-repeats may be a result of interactions with different binding partners, or alternatively may regulate the interaction between this region of the lncRNA and its cognate RBPs.

### The lncRNA SLNCR1

The lncRNA SLNCR is associated with the genesis of melanoma cells^64,65^. The primary isoform (SLNCR1) upregulates matrix metalloproteinase-9 (MMP-9) through interactions with androgen receptor (AR) and brain-specific homeobox protein 3a (Brn3a)^64^. Only a single highly conserved region of SLNCR1 – between nts 372 – 672 – is needed for increased melanoma invasion^23^. The Brn3a and AR binding sites on SLNCR1 are directly adjacent to each other, at nts 462 – 572 and 610 – 637, respectively^23^. AR favors binding single-stranded regions of RNA: AR binding is stifled in the presence of RNA secondary structure^23^. Mutating the AR binding sites on SLNCR1 attenuated melanoma invasion, suggesting the interaction between SLNCR1 and AR is critical for the carcinogenic properties of SLNCR1^23^.

The conserved region of SLNCR1 (nts 372 – 672) was reported to be unstructured from both *in-cell* and *in vitro* SHAPE– and DMS-MaP experiments, possessing high median reactivities and Shannon entropies^23^. Particularly, the Brn3a binding site was found to be completely unstructured^23^. However, the AR binding motifs possessed only moderate reactivities and Shannon entropies, suggesting the presence of multiple distinct paired states^23^. Although assumed to be unstructured, arcplot-supported models of the region of SLNCR1 harboring the AR binding sites were reported based on the DMS-MaP data^23^. It is possible that SLNCR1 only transiently samples a structured state. We hypothesize that SLNCR1 alternates between multiple states, potentially modulating its interaction with AR by sampling both structured, AR-incompetent, and unstructured, AR-competent, conformations.

To understand whether the conserved region of SLNCR1 samples different structured states, we used DRACO to deconvolute the Schmidt *in vitro* DMS-MaP dataset^23^. DRACO identified one large region (nts 600 – 780; **Figure 2A**) that spans both the AR binding motifs (nts 612 – 620, and 627 – 635), but not the Brn3a binding site (nts 462 – 572). This region was reported by DRACO to sample two distinct folds in a 0.64: 0.36 stoichiometry (**Figure 2B**).

**Figure 2.**
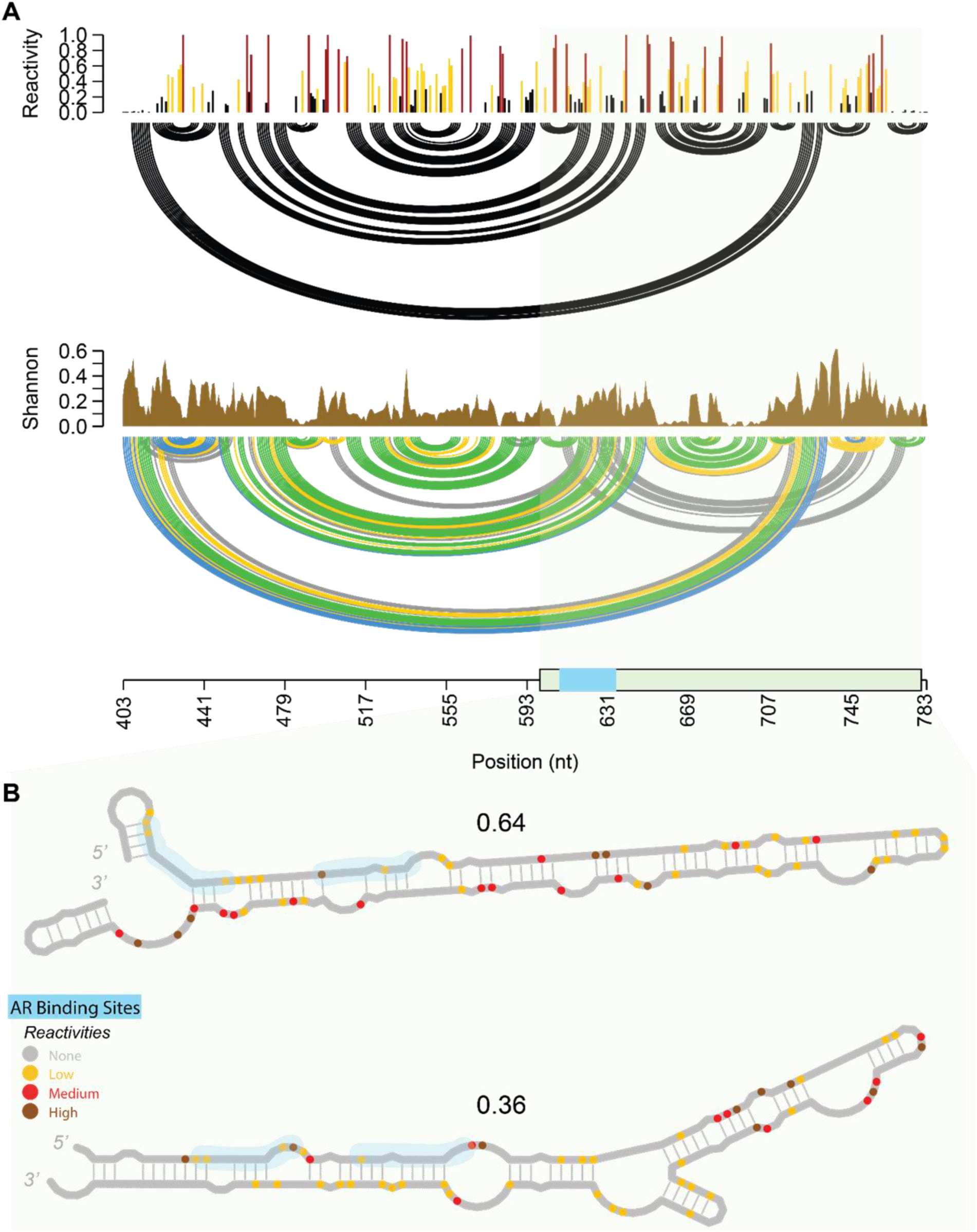
Deconvolution of the human SLNCR1 lncRNA from the Schmidt chemical probing dataset^23^. **A** The reactivity profile, arc plot, and Shannon entropies for the conserved region of SLNCR1. The green sausage plot represents the region identified by DRACO to sample multiple conformations. The blue region in the sausage plot represents the AR binding sites. **B** The two deconvoluted SLNCR1 conformations are shown, along with their respective populations reported by DRACO. The AR binding sites are highlighted in blue.

The deconvoluted minor fold of SLNCR1 is congruent with the arcplot-supported structure reported in the SLNCR1 chemical probing paper^23^. Both the deconvoluted minor fold, and the published arcplot-supported structure are composed of a long stem punctuated by five internal loops, with the third internal loop harboring a short stem capped by a “GGUU” tetraloop^23^. The minor deconvoluted structure and arcplot-supported structure possess a “UUCCU” pentaloop at the apical region of the long stem (**Figure 2B**). The deconvoluted major fold of SLNCR1 identified by DRACO is distinct from the arcplot-supported structure reported in the Schmidt *et al.* paper^23^. This major fold consists of one large stem loop with two smaller stem loops flanking it on the 5’ and 3’ sides (**Figure 2B**). Strikingly, the first AR binding motif is remodeled in this dominant structure compared to the minor structure, being split between the smaller 5’ hairpin and the large stem. The second AR binding motif is largely paired and unchanged in the two deconvoluted structures: possessing four unpaired nts in the major conformation compared to only one unpaired nt in the minor conformation. That the two AR binding motifs are both paired in the two deconvoluted conformations is surprising, as AR has been shown to preferentially bind unstructured RNAs^23^. The presence of two structured folds in the AR binding motifs suggests that SLNCR1 may adopt different structures to regulate its interactions with AR, facilitating binding only in an unpaired state. It is unclear if either of these conformations would facilitate AR binding, or if this region must be completely unstructured for AR binding. Future studies will experimentally validate the presence of these different structured states, and their relationship to AR binding.

The fact that DRACO could recover a previously reported structure from the DMS-MaP data inspires confidence in its ability to accurately deconvolute RNA conformations. Further, the Brn3a binding site was originally reported as unstructured, possessing high SHAPE reactivities and Shannon entropies^23^. In agreement with this, DRACO did not identify this region as harboring coexisting conformations. These results act as a form of validation for DRACO – ensuring that (i) DRACO can recover known folds from DMS-MaP data, and (ii) RNA regions known to be unstructured are not identified by DRACO to harbor coexisting folds.

### The lncRNA GAS5

The lncRNA GAS5 is multifunctional: it both acts as a micro-RNA (miRNA) sponge for miRNA-21 and miRNA-135, and binds nuclear steroid receptors (SR) to induce apoptosis^66–69^. GAS5 functions as a tumor suppressor, and its misregulation is implicated in many types of cancers, including breast cancer, prostate cancer, and esophageal cancer^70,71^.

The structure of GAS5 was characterized in a 2020 study by SHAPE-MaP both *in vitro* and *in cellulo*^22^. GAS5 was reported to fold in a modular fashion with three structured regions: a 5’ module, a core module, and an SR binding domain (**Figure 3A**). The 5’ module (nts 1 – 170) was reported as only mildly structured, possessing long stretches of single-stranded RNA, as this region binds miRNA-21 and miRNA-135, and is believed to interact with proteins regulating the cell cycle^22,69,72^. The 5’ domain possesses low-to-moderate SHAPE reactivities, and moderate Shannon entropies, indicative of a structured region sampling different base pairing patterns (**Figure 3A**).

**Figure 3.**
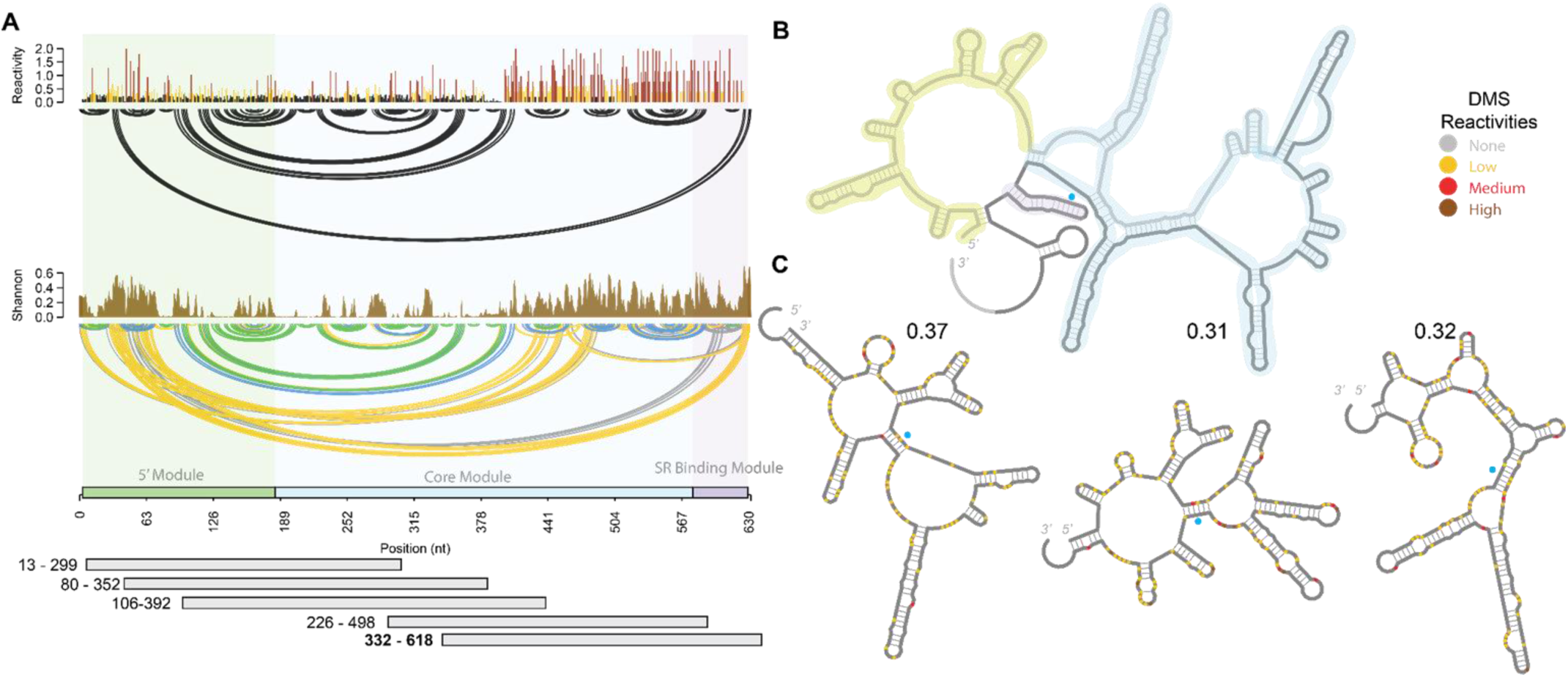
Deconvolution of the human GAS5 lncRNA by DRACO. **A** Arc plot showing the reactivities and Shannon entropy values of GAS5 in the original Frank et al chemical probing dataset^22^. The different domains of GAS5 are colored in green, blue, and purple, respectively. The horizontal grey bars represent a region identified by DRACO to possess coexisting conformations. **B** The published structure of GAS5^22^. **C** The structures of three coexisting folds identified by DRACO in one of the deconvoluted regions (nts 332 – 618) of GAS5 are shown along with their calculated stoichiometries. The colored points on the deconvoluted RNA structures represent 1M7 reactivities, with yellow as mildly reactive, red as moderately reactive, and purple as highly reactive. The blue dot on each structure indicates G549.

The core module of GAS5 (nts 172 – 540) is highly structured, possessing primarily low SHAPE reactivities and low Shannon entropies (**Figure 3A**). This region helps mediate mammalian target of rapamycin (mTOR) signaling pathways^22^. The core module was proposed to adopt a structured tertiary structure, potentially acting as a scaffold to engage RBPs^22^. Overexpressing either the 5’ module or central core module amplifies apoptosis^22^. The larger SHAPE reactivities and Shannon entropies towards the 3’ end of the core module suggest some part of this region may sample multiple base pairing patterns (**Figure 3A**). Immunoprecipitation assays of individual modules of GAS5 show *hundreds* of proteins that bind GAS5 when pulled down (448 and 619 proteins to the 5’ and core modules, respectively), implying overlap between RBP binding regions^22^.

Lastly, the SR binding domain (nts 541 – 573) helps regulate the lifecycle of cells dependent on steroids for cellular growth. Notably, mutating a single nucleotide (G549A) in the SR binding module abolishes SR binding and arrests cell growth^73^. We hypothesize that GAS5 alternates between different conformations, either due to RBP binding events, or spontaneously in order to be recognized by different RBPs.

We employed DRACO to deconvolute the Frank *et al. in vitro* chemical probing dataset used to characterize the structure of GAS5^22^. DRACO identified five overlapping regions in GAS5 with coexisting conformations, which combined span the entire length of the lncRNA (**Figure 3A**). Typically, overlapping windows with multiple conformations are merged in DRACO if the overlapping windows possess the same number of coexisting conformations^25^. However, each region of GAS5 deconvoluted by DRACO was found to harbor a different number of coexisting conformations, creating multiple overlapping windows of conformational variability. None of the regions of conformational variability identified by DRACO lie in a single domain of GAS5, suggesting that the modular, three-domained fold of GAS5 identified by Frank *et al.* may not be the only structure this lncRNA samples^22^.

One of the regions identified by DRACO, between nts 332 – 618, spans both the core module and SR binding domain of GAS5 (**Figure 3B**). The deconvoluted structures for this region reported by DRACO completely remodel the core module and SR binding domain, and were reported to exist in near equal populations. Inspection of G549 – a nucleotide in the SR binding domain whose mutation abolishes SR binding – in the deconvoluted structures reveals this nucleotide lies in a dynamic region. The characteristic hairpin of the SR binding domain originally reported is reconfigured into a highly structured stem flanked by two junctions on the 5’ and 3’ sides in all three deconvoluted structures^22^.

Mutation of G549 could potentially disrupt or destabilize this stem linking the two junctions, possibly distorting the RNA structure, and explaining how mutation of this single nucleotide abolishes SR binding. As a qualitative test on the importance of this single nucleotide in the overall fold of this region of GAS5, we used MFold to predict the structure of this region with either G549 or the G549A mutation^74^. Indeed, mutating this single nucleotide drastically changes the structure of this region of GAS5 reported by MFold (**Supplementary** Figure 2)^74^. This suggests that the SR binding domain in GAS5 does not necessarily exist as a single, modular domain – but instead can exist in multiple structured states. The conformational variability of the SR binding domain may allow GAS5 to regulate its interactions with SR.

The 5’ module of GAS5 was originally reported to be only moderately structured, possessing multiple short stretches of paired nucleotides^22^. The binding sites for miRNA-21 and miRNA-135 both lie in the 5’ module^22,69^. The relative lack of structure in this region is consistent with RNA binding activity and efficient translation initiation. Interestingly, DRACO reported these regions as structured, participating in long stems in the deconvoluted structures (**Supplementary** Figure 2). Both deconvoluted structures of the 5’ module of GAS5 hold multiple bulges and internal loops, suggesting the long stem structures may spontaneously reorganize.

GAS5 was originally reported to fold in modular domains, with specific binding partners and functions identified for each region^22^. The multi-functionality of GAS5 was attributed to the presence of multiple structured elements throughout the different domains. However, the identification of hundreds of RBPs binding GAS5 in RNA-pulldown experiments suggests the GAS5 lncRNA may sample multiple different structured states. Our analysis with DRACO suggests GAS5 is conformationally dynamic, with each region sampling multiple distinct geometries. This lncRNA may adopt different conformations to modulate its interactions with binding partners, and by extension its specific cellular function.

### The lncRNA MALAT1

The lncRNA MALAT1 is a highly conserved and relatively well-studied ∼7kb lncRNA that helps regulate metastasis^75–78^. It was first identified in non-small cell lung cancer cells, and is associated with a large number of disease states and cancer types^75,76,79^. MALAT1 is highly expressed in a broad range of tissue types, but is retained in the nucleus, and often localizes to nuclear speckles^7,76^. Consequently, MALAT1 has been shown to associate with many proteins that localize to nuclear speckles: RNA-binding protein with serine-rich domain 1 (RNPS1), serine/arginine repetitive matrix protein 1 (SRm160), and intron binding protein 160 (IBP160) are essential for MALAT1 localization to nuclear speckles. And serine and arginine rich splicing factor 1 (SRSF1), SON1, hnRNPC, and hnRNPH1 have also been observed to bind MALAT1^80–83^. The interactions between MALAT1 and RBPs contribute to its role in regulating pathways around cellular metastasis.

Two different structures have been proposed for the entire MALAT1 transcript. A computational study proposed a secondary structural model by calculating the minimum free energy structure using previously-published in-cell DMS-MaPseq data^84^. A more recent study reported the secondary structure of MALAT1 through chemical probing using 5-nitroisatoic anhydride (5NIA) both *in vitro* and in cell^21^. The latter study reported very few differences in the structure of MALAT1 inside and outside the cellular environment, suggesting that cellular crowding / RBPs have little effect on the fold of MALAT1. The experimentally reported structure of MALAT1 is composed of small, modular base-paired regions^21^. The 3’ end of MALAT1 lacks a canonical poly-A tail, and instead holds a highly conserved and exceedingly stable triple helix that protects the RNA from 3’-exonucleolytic degradation^85,86^.

We used DRACO to deconvolute the *in vitro* Munroy-Eklund chemical probing data^21^. DRACO identified three regions of conformational variability, each spanning approximately ∼250 nts (**Figure 4**). Each of these three regions of conformational variability possess moderate to high reactivities, but only moderate Shannon entropies (**Figure 4A**). The first region of conformational variability identified by DRACO (nts 1,562 – 1,823) harbors two conformations that exist in a 0.62: 0.38 ratio. The other two regions (nts 5,088 – 5,349 and 7,001 – 7,262) alternate between dominant, more linear structures, and minor, more modular structures (**Figure 4B**), with a population ratio of 0.94: 0.06.

**Figure 4.**
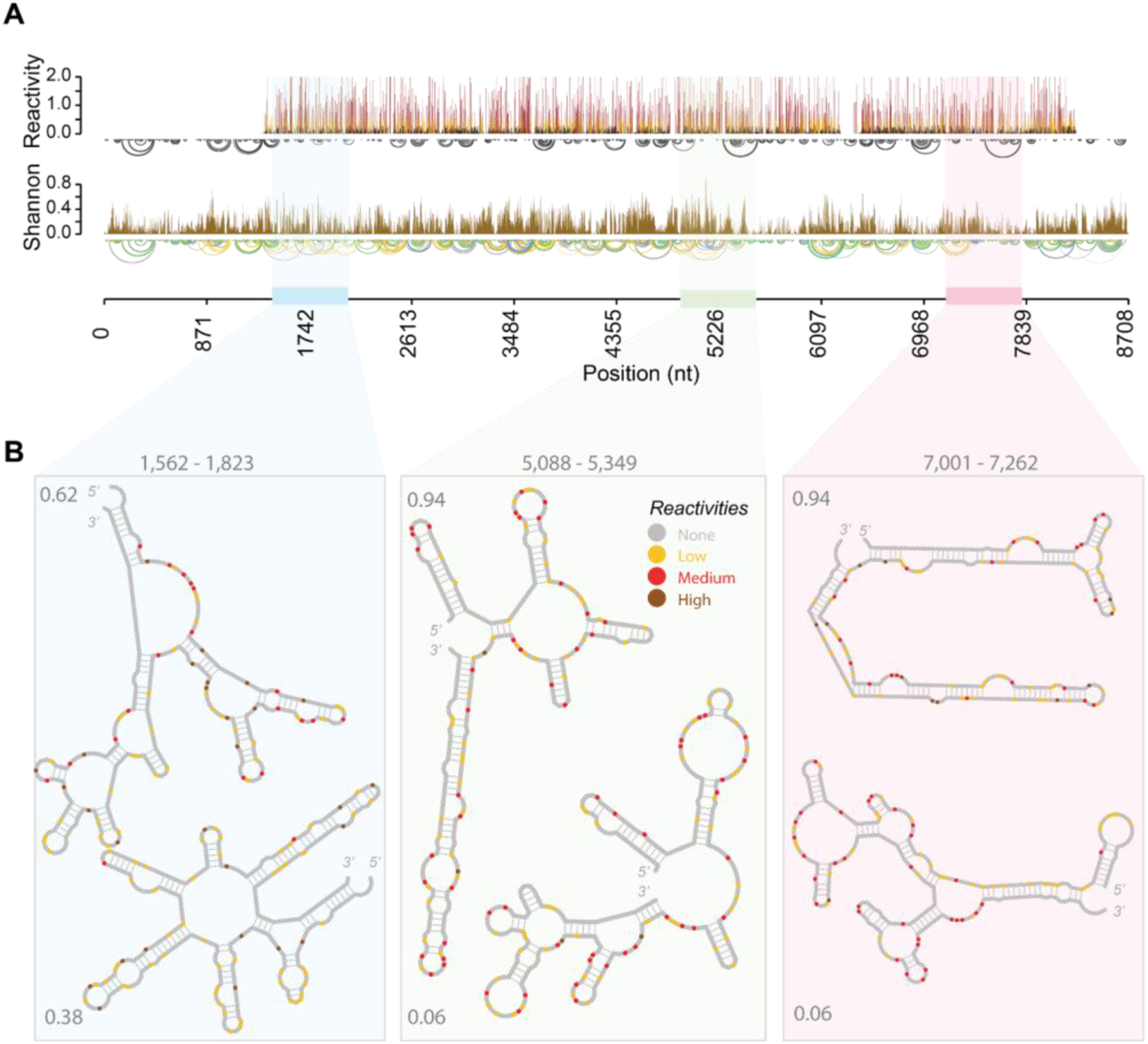
Deconvolution of the human MALAT1 lncRNA using DRACO. **A** Arc plot and deconvoluted reactivities and Shannon entropies reported by deconvoluting the Munroy-Ekland chemical probing dataset^21^. The blue, green, and pink rectangles represent the conformationally dynamic regions identified by DRACO. **B** The three regions of conformational variability with their respective secondary structures, deconvoluted reactivities, and stoichiometries as reported by DRACO.

DRACO can identify alternate structures with a population threshold of 5%, thus the two minor conformations with a 6% population ratio are potentially less confident than the minor geometry in the first region (nts 1,562 – 1,823)^25^. However, the final two regions of conformational variability identified by DRACO (nts 5,088 – 5,349 and 7,001 – 7,262) possess moderate-to-high reactivities, and low-to-moderate Shannon entropies, suggesting these regions possess multiple paired states. With the general topology and base pairing of these conformationally dynamic regions deconvoluted by DRACO, future experiments – such as SAXS or NMR – may be done to identify whether these regions of MALAT1 do indeed sample multiple structured states.

The lncRNA MALAT1 has been shown to interact with a large number of binding partners, including proteins, U1 small nuclear RNA, and different miRNAs^84,87^. However, the exact binding sites on MALAT1 between many of its numerous binding partners have not been extensively characterized. Future work will seek to characterize the exact binding locations on MALAT1 of its different binding partners, and whether these binding events are associated with the conformationally dynamic regions identified here, and specific conformational states.

### The lncRNA MEG3

The lncRNA MEG3 possesses multiple splice variants and is located primarily in the nucleus. It induces apoptosis by stimulating the p53 pathway, and is essential for neuronal development^88,89^. Work by Zhu *et al.* found that MEG3 binds the protein clusterin, which is overexpressed in several types of cancer^90,91^. Further, the expression of MEG3 has been observed to inhibit tumor proliferation, suggesting MEG3 has anti-carcinogenic effects – possibly by inhibiting Clusterin transcription^92,93^.

The structure of MEG3 has been solved by SHAPE– and DMS-MaP^24^. The most common splice variant of MEG3 is 1,595 nts and possesses five structural domains that reportedly fold modularly within exon boundaries^24^. Exon 3 is the most evolutionarily conserved region of MEG3, and reportedly folds into two highly stable structural domains (D2 and D3; **Figure 5B**)^24^. Exon 3 alone was shown to stimulate the p53 pathway, and deletion of either D2 or D3 in exon 3 abolished MEG3-induced stimulation of the p53 pathway. Deletion or mutation of a specific hairpin (H11) in D2 markedly stifled activation of the p53 pathway^24^. This H11 hairpin in the D2 domain is hypothesized to form long-range interactions with a separate hairpin in the D3 domain, due to evolutionarily conserved complementary sequences that would potentially allow for long-range kissing hairpin interactions^24^. However, the exact mechanism by which the D2 and D3 domains stimulate the p53 pathway are unknown.

**Figure 5.**
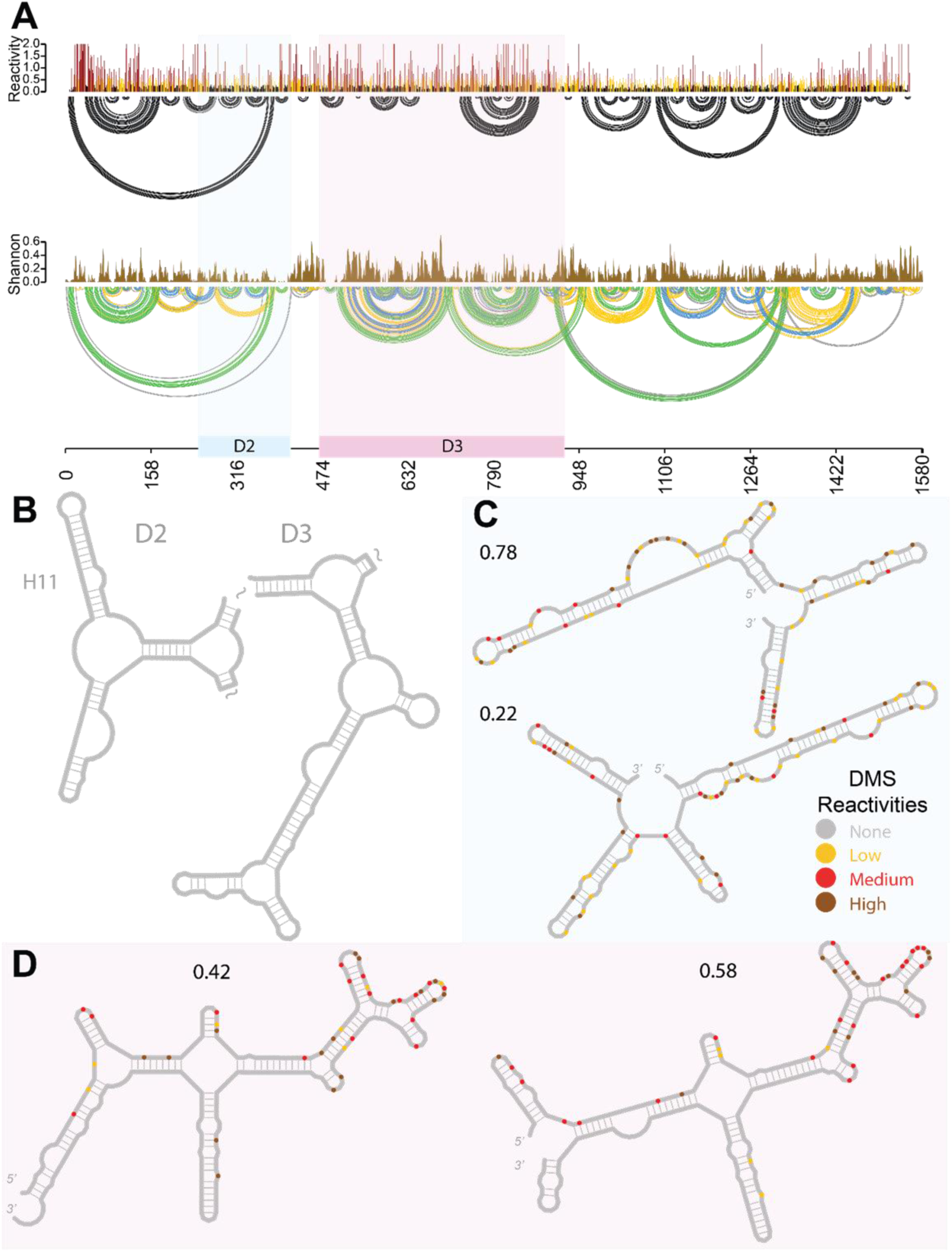
Deconvolution for the human lncRNA MEG3 using the Uroda chemical probing dataset^24^. **A** Arc-plot, DMS reactivities, and Shannon entropy for MEG3 using the Uroda chemical probing dataset^24^. The conserved D2 region is highlighted in blue, and the D3 region in pink. **B** The experimentally reported structure of the D2 and D3 domains of MEG3. **C, D** Coexisting structures in the D2 and D3 regions, respectively, with their stoichiometries deconvoluted by DRACO, colored according to the deconvoluted DMS reactivities.

We used DRACO to deconvolute the Uroda chemical probing data to explore whether the functionally significant regions of MEG3 are structurally dynamic^24^. We deconvoluted both the 1M7 and DMS-MaP datasets separately to identify whether any coexisting conformations are reproduced across the different probing reagents. DRACO identified two regions of conformational variability present in both chemical probing datasets. One of these regions (nts 153 – 373) lies in the structured D2 domain in the evolutionarily conserved exon 3 (**Figure 5D**). The two alternate conformations identified by DRACO retain the H11 hairpin loop, but show a substantial reorganization of base pairing patterns upstream of the conserved H11 hairpin, suggesting the two hairpin loops may be conformationally static. Conversely, the clusterin binding site on MEG3 was previously reported between nts 732 – 1174 based on electrophoretic mobility shift assays on different MEG3 fragments^90^. This clusterin binding region overlaps with the conformationally dynamic region of D3 identified in this study (397 – 653), suggesting that the interactions of MEG3 with clusterin are potentially modulated by or modulate the conformation of the D3 domain of MEG3. Future work will identify the exact binding site between clusterin and MEG3, and explore whether clusterin binding is sensitive to different conformations of this region of MEG3.

### The lncRNA NEAT1

The NEAT1 (nuclear enriched abundant transcript 1) lncRNA possesses two isoforms: a short 3.7 kb isoform (NEAT1_S), and a long ∼23 kb isoform (NEAT1_L). Both of these lncRNAs are involved in nuclear paraspeckle assembly, however, only the longer isoform (NEAT1_L) is required for the formation of nuclear paraspeckles^7,94^.

A single structure has been reported for the NEAT1_S lncRNA using SHAPE-seq, which exhibits a large degree of order with different structured domains^95^. To our knowledge, no structure has been reported for the larger isoform NEAT1_L, which contains the NEAT1_S isoform at the 5’ end – implying a degree of structural conservation between the two different sized transcripts.

To understand whether any areas of NEAT1 harbor coexisting folds, we used DRACO to deconvolute the in cell Rouskin chemical probing dataset^25,96^. We could not use the Lin *et al.* chemical probing dataset that originally reported the structure of NEAT1_S for structural deconvolution, as DRACO was designed only for the deconvolution of chemical probing experiments done using *mutational* profiling.

DRACO performed exactly as intended: it identified 94 regions on NEAT1_L predicted to sample multiple conformations. However, mapping the reads from in cell DMS-MaP experiments performed on the entire transcriptome against the NEAT1 representative transcript resulted in spurious reads, causing an overestimation of the number of reads corresponding to NEAT1, and complicating structural deconvolution by DRACO. In reality, the number of reads from the in cell DMS-MaP dataset that map to NEAT1 are much lower. DRACO treated reads that were falsely mapped to NEAT1 as part of the data to be deconvoluted, yielding an inaccurate description of conformationally dynamic regions in NEAT1_L. This is reflective of several issues we faced when trying to deconvolute in cell SHAPE– / DMS-MaP datasets performed on the entire transcriptome, which we discuss in the next section.

### The Struggles Around Deconvoluting lncRNA Structural Dynamics with In-Cell Chemical Probing Data for Low Abundance RNAs

The lncRNA structures reported here describe coexisting conformations of lncRNAs *in vitro*. The conformational ensemble a lncRNA samples may be different *in vitro* versus in the cell. At the start of this project, we attempted to identify coexisting lncRNA conformations by deconvoluting in cell chemical probing data, but we quickly encountered obstacles that made this unfeasible. We think it is beneficial for the community to discuss these here.

We attempted to identify coexisting conformations of 12 lncRNAs by deconvoluting entire in-cell DMS-MaP datasets (**Supplementary Table 2**). There are two (human) in-cell DMS-MaP datasets with enough sequencing depth to potentially facilitate structural deconvolution of lncRNAs^96,97^. We first mapped the in-cell reads to the canonical (most abundant) lncRNA reference transcripts. However, mapping in-cell DMS-MaP datasets to individual transcripts using Bowtie2 resulted in spurious reads, greatly overestimating the sequencing depth for each lncRNA (**Supplementary** Figure 3)^98^. In a word, considering that lncRNAs possess several isoforms and that reads could correspond to *any* of these, it is not appropriate to map genome-wide reads to a single reference transcript. This is especially true for short reads obtained via Illumina Platform sequencing approaches.

We then attempted to map the in-cell DMS-MaP datasets to the entire transcriptome, using STAR with several different genome annotations (e.g. MANE select, lncRNA annotation, ENSEMBL)^99^. Regardless of the annotation used, mapping to the transcriptome resulted in an *under*estimate of sequencing depth for each of the 20 lncRNAs of interest. The lncRNA transcripts present in cells at the time of chemical probing may not correspond to the annotated transcripts. This could be due to alternative splice forms in the cells.

Additionally, the typical read-length in mutational profiling experiments is not long enough to map to specific lncRNA spliceforms. The reads from the in-cell datasets are typically 100 – 200 nts long^96,97^. However, the shortest lncRNA examined in this study is ∼650 nts long. Multiple different splice forms of the same lncRNA may be present in a cell at any given time. Even for the shortest lncRNA studied here, identifying which splice form each read maps to is challenging, if not impossible, due to the short read-length. The inability to to map individual reads to specific splice forms of a lncRNA further obscures and complicates the structural deconvolution of lncRNAs. It is possible that a lncRNA primarily samples one structure in one splice form, and different structure when in another splice form. However, the short read length utilized in most chemical probing experiments prevents the assignment of individual conformations to specific splice forms using structural deconvolution methods. Additionally, the window size in DRACO is by default set equal to 90% of the median read length, which intrinsically limits the size of the RNA region being deconvoluted. Hypothetically, a lncRNA could alternate between two conformations that reconfigure its entire structure, but DRACO would only be able to deconvolute *local* regions of the RNA that sample multiple conformations. This was observed with our initial investigation of the dynamics of lncRNA structures in cells (**Supplementary Figures S4, S5, S6**).

Finally, the relatively low-abundance of lncRNAs complicates the ability to attain a sufficient number of reads for structural deconvolution. Many lncRNAs are expressed in low concentrations, or their expression levels are strictly temporally regulated^100^. The lncRNA Paupar was reported to have ∼15 copies per cell measured by chromatin isolation by RNA purification (ChIRP) assays^101^. Consequently, the number of reads needed to achieve a sequencing depth sufficient for the structural deconvolution of low-abundance lncRNAs is unfeasible. Deconvolution algorithms like DRACO require a sequencing depth of at least ∼5,000 to identify coexisting conformations^25^. Achieving a sequencing depth of 5,000 for *every* RNA transcript in a cell – particularly the low-abundance lncRNAs – would require billions to trillions of reads.

We concluded that deconvoluting coexisting lncRNA structures from in-cell SHAPE– / DMS-MaP datasets is challenging because of (i) overestimation of sequencing coverage when mapping in-cell chemical probing to individual lncRNA transcripts, (ii) underestimation of sequencing coverage when mapping to the transcriptome because of asynchronous annotations, and (iii) average read-lengths being too short. Future in-cell SHAPE– / DMS-MaP datasets analyzed using long-read sequencing approaches (eg PACBIO or Oxford Nanopore) may circumvent these issues as the technology and protocols continue to develop and evolve^16,102,103^. However, for now structural deconvolution of lncRNAs using DRACO works best on *individual* SHAPE– / DMS-MaP datasets conducted either *in vitro* or *in cell* where the lncRNA of interest is either expressed in high concentrations, or is the sole species being sequenced.

## Conclusion

The secondary structures of many RNAs are deduced using reactivity profiles from chemical probing experiments. However, the resulting reactivity profiles do not represent a single structure, but instead represent the *average* reactivity profile of all RNA conformations present at the time of chemical probing. Many RNAs are structurally dynamic, and adopt multiple different conformations in different populations. Thus, the secondary structures for RNA transcripts solved by chemical probing experiments are often *average* structures. Computational methods, such as DRACO, DREEM, and DANCE-MaP, deconvolute the reactivity profiles from chemical probing experiments and report coexisting conformations whose individual reactivity profiles *average out* to the observed reactivity profile^25,31,32^.

We sought to identify coexisting conformations of five lncRNA transcripts by deconvoluting publicly available *in vitro* or in-cell SHAPE– / DMS-MaP datasets. We used DRACO to deconvolute the structural ensembles of the lncRNAs XIST, SLNCR1, GAS5, MALAT1, and MEG3^6,21–24^. We found that each of these lncRNAs possesses at least one conformationally variable region, many of which have been previously reported as sites of RBP interactions. The conformational variability of these lncRNAs may contribute to their multifunctionality: lncRNAs might spontaneously sample different conformations in response to RBP binding events, or in order to regulate their interactions with RBPs. Future studies will validate the presence of these coexisting geometries using SAXS, NMR, and mutational experiments – and whether these coexisting geometries are functionally relevant. We also discuss the limitations with deconvoluting in cell SHAPE– / DMS-MaP experiments, in the hopes the field will continue to evolve to eventually facilitate the structural deconvolution of lncRNAs throughout the entire transcriptome.

## Acknowledgements

We would like to thank David Klingler and Daniel Cohn for HIV-1 RRE chemical probing data, Danny Incarnato for useful conversations, and the rest of the Jones lab for comments and feedback.

## Conflicts of Interest

A.N.J. is a consultant for NextRNA.

## Funding

This work was funded by the National Science Foundation, 2243667 to A.N.J.

