## Supplemental Information for "Alternative Conformations of lncRNAs Identified Through Structural Deconvolution of SHAPE- and DMS-MaP Datasets"

### Supplementary Information

**Supplementary Table 1.** List of the lncRNAs studied in this report and their corresponding length, gene ID, the transcript ID used in this study, the dataset that was deconvoluted using DRACO, and the number of reads used for deconvolution.

| lncRNA | Length (nts) | Ensembl Gene ID | Transcript ID | Chemical Probing Data Used | Number of Reads |
|---|---|---|---|---|---|
| XIST (Mouse) | 17,769 | ENSMUSG00000086503.3 | ENSMUST00000127786.2 | Smola *et al.*[6] | 812,308 |
| SLNCR1 | 2,527 | ENSG00000227036.11 | ENST00000649350.1 | Schmidt *et al.*[23] | 10,736,792 |
| GAS5 | 632 | ENSG00000234741 | ENST00000450589.5 | Frank *et al.*[22] | 4,780,514 |
| MALAT1 | 8,708 | ENSG00000251562 | ENST00000534336.1 | Monroy-Eklund *et al.*[21] | 4,733,945 |
| MEG3 | 1,582 | ENSG00000214548 | ENST00000451743.6 | Uroda *et al.*[24] | 1,428,645 |

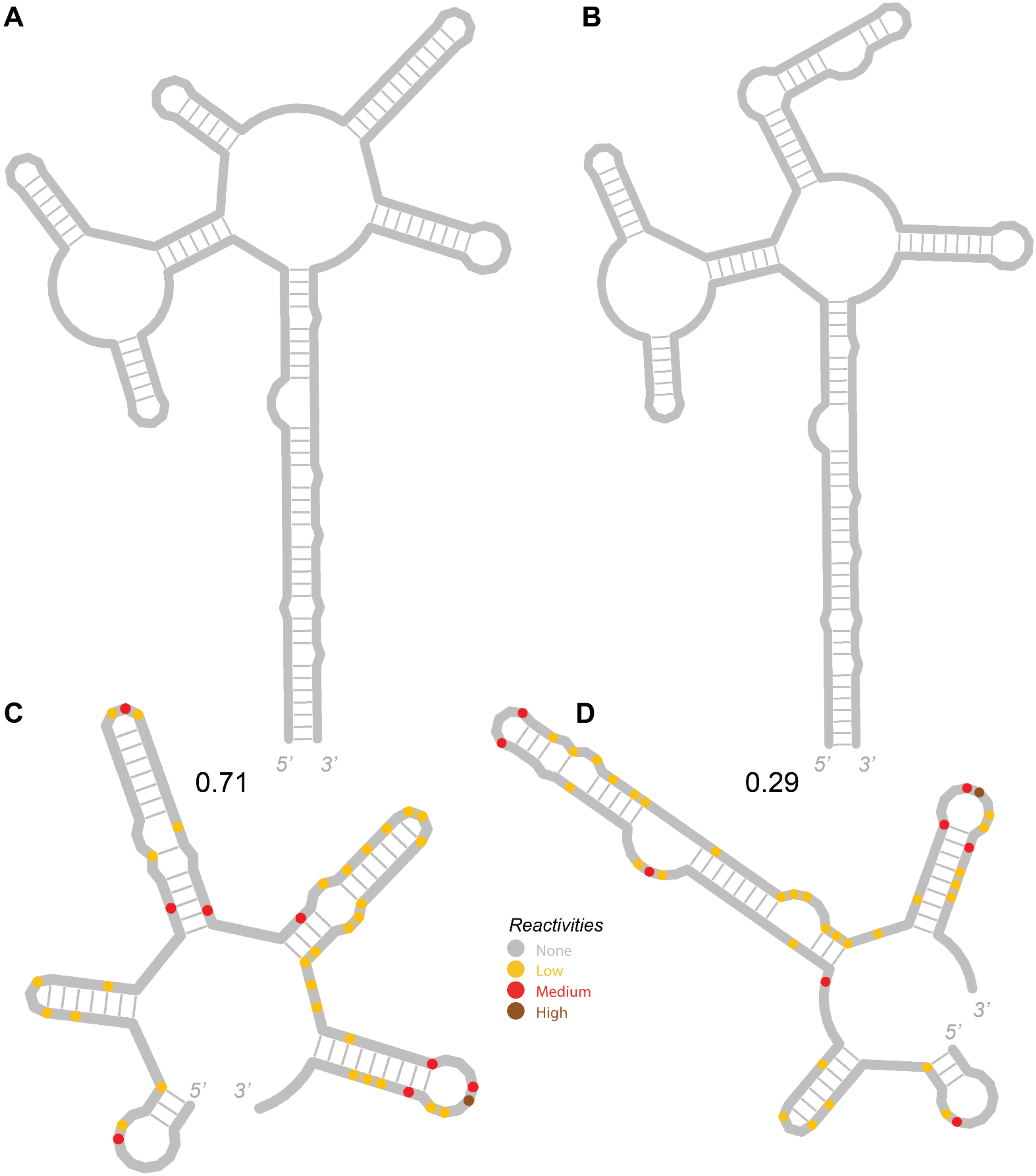


**Supplementary Figure 1.** Validating the ability of DRACO to deconvolute SHAPE-MaP data using known folds of the HIV RRE. **A.** The dominant, 5-SL structure of the RRE, as reported by Kjems *et al.*[104]. **B.** The alternate, 4-SL structure of the RRE, as reported by Sherpa *et al.*[105]. **C, D** Coexisting folds of the HIV RRE recovered by DRACO by deconvoluting SHAPE-MaP chemical probing data. DRACO was able to identify the 5-SL and 4-SL structures, along with their reported stoichiometries. The nts are colored by their normalized reactivities.

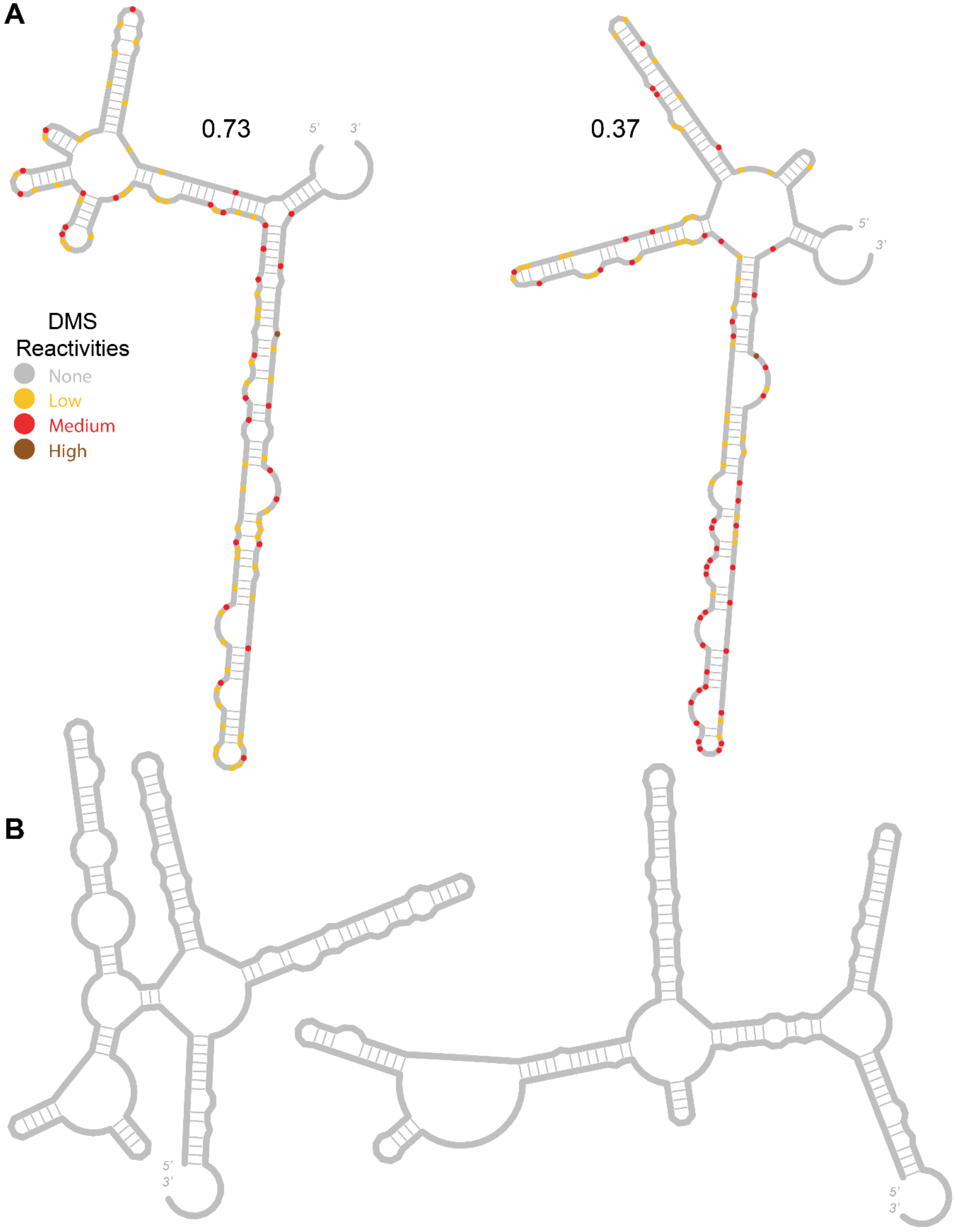


**Supplementary Figure 2.** Additional analysis on the lncRNA GAS5. **A.** Coexisting structures in the lncRNA GAS5 identified by DRACO lying in the 5' module, between nts 13 - 299. The individual nts are colored according to their deconvoluted reactivities. **B.** MFold-predicted structures of the region of GAS5 deconvoluted by DRACO discussed in main text (nts 332 - 618), with the nascent sequence on the left and the G549A mutated sequence on the right.

**Supplementary Table 2.** List of the lncRNAs we attempted to deconvolute using entire transcriptome in cell SHAPE- / DMS-MaP data. Listed are the lncRNA names, published secondary structures, the Ensembl Gene ID of the gene, and the Transcript ID of each lncRNA studied.

| lncRNA | Published Secondary Structure | Ensembl Gene ID | Transcript ID |
|---|---|---|---|
| XIST | Wutz 2002<br>Duszczyk 2008<br>Maenner 2010<br>Duszczyk 2011<br>Fang 2015<br>Weeks 2016<br>Ponti 2016<br>Lu 2016<br>Smola 2016<br>Rivas 2016<br>Liu 2017 | ENSG00000229807 | ENST00000429829.5 |
| JPX | Karner 2020 | ENSG00000225470 | ENST00000639185.1 |
| SLNCR1 | Novina 2020 | ENSG00000227036 | ENST00000568947.2 |
| MALAT1 | Lin 2018 | ENSG00000251562 | ENST00000534336.1 |
| GAS5 | Ortlund 2020 | ENSG00000270084 | ENST00000430245.5 |
| NEAT1 | Nakagawa 2019 | ENSG00000245532 | ENST00000499732.2 |
| TCF7 | Somarowthu 2019 | ENSG00000081059 | ENST00000522653.5 |
| FTX | N/A | ENSG00000230590 | ENST00000602812.5 |
| FIRRE | N/A | ENSG00000213468 | ENSG00000213468.4 |
| INE1 | N/A | ENSG00000224975 | ENST00000456273.1 |

| | | | |
|---|---|---|---|
| ZNF674-AS1 | N/A | ENSG00000230844 | ENST00000421685.2 |
| PANDAR | N/A | ENSG00000281450 | ENST00000629595.1 |

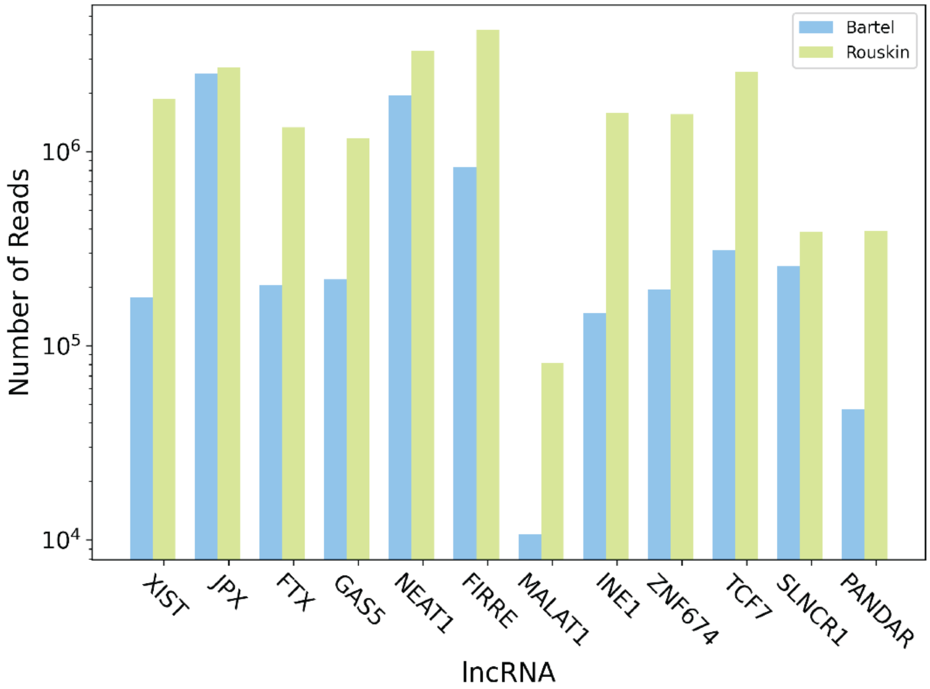


**Supplementary Figure 3.** The spurious coverages reported by attempting to map and deconvolute the conformational ensembles of 12 lncRNAs using two different in-cell chemical probing datasets done against the entire transcriptome. The blue and green bars represent the sequencing depth for the Bartel and Rouskin chemical probing datasets, respectively.

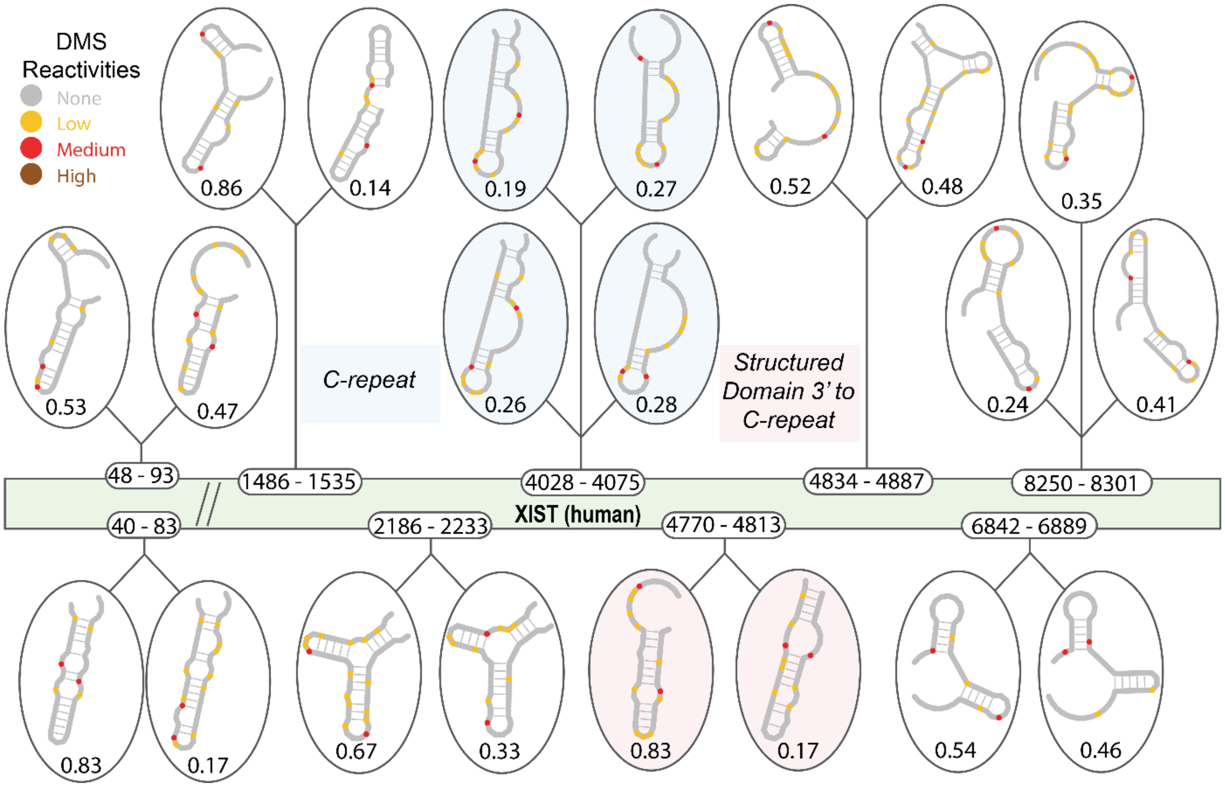


**Supplementary Figure 4.** Attempted structural deconvolution of lncRNA XIST (human) using in cell SHAPE-MaP data against the entire transcriptome. The short window size of DRACO, stemming from the short read lengths of the chemical probing experiments, led to the spurious identification of small, local structurally heterogeneous regions.

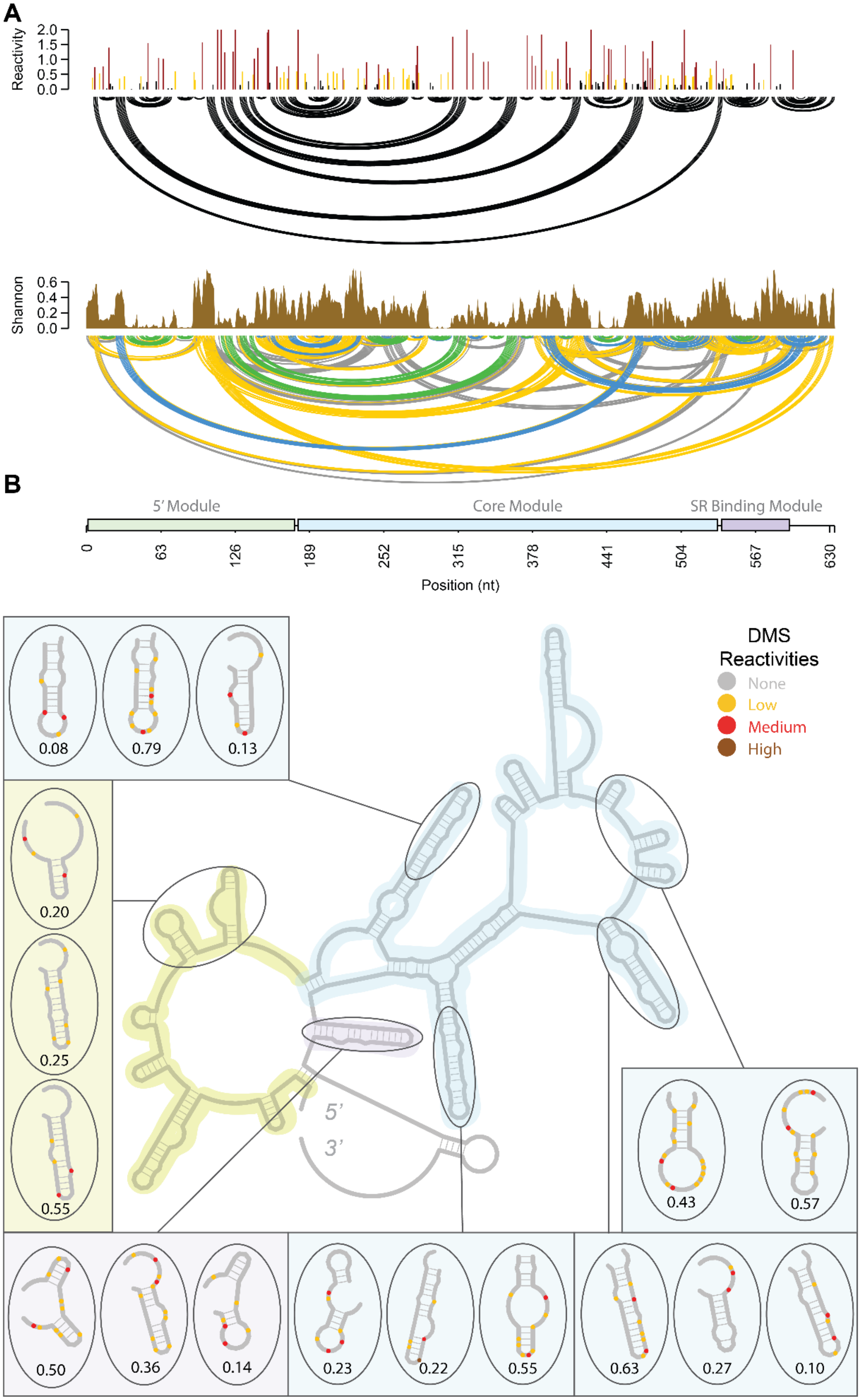


**Supplementary Figure 5.** Attempted structural deconvolution of lncRNA GAS5 using in cell SHAPE-MaP data against the entire transcriptome. The short window size of DRACO, stemming from the short read lengths of the chemical probing experiments, led to the spurious identification of small, local structurally heterogeneous regions. **A.** The reactivities, arc plot, and Shannon entropies for GAS5. **B.** The published structure of GAS5, and the regions of conformational variability spuriously identified by DRACO from the in cell SHAPE-MaP dataset done against the entire transcriptome.

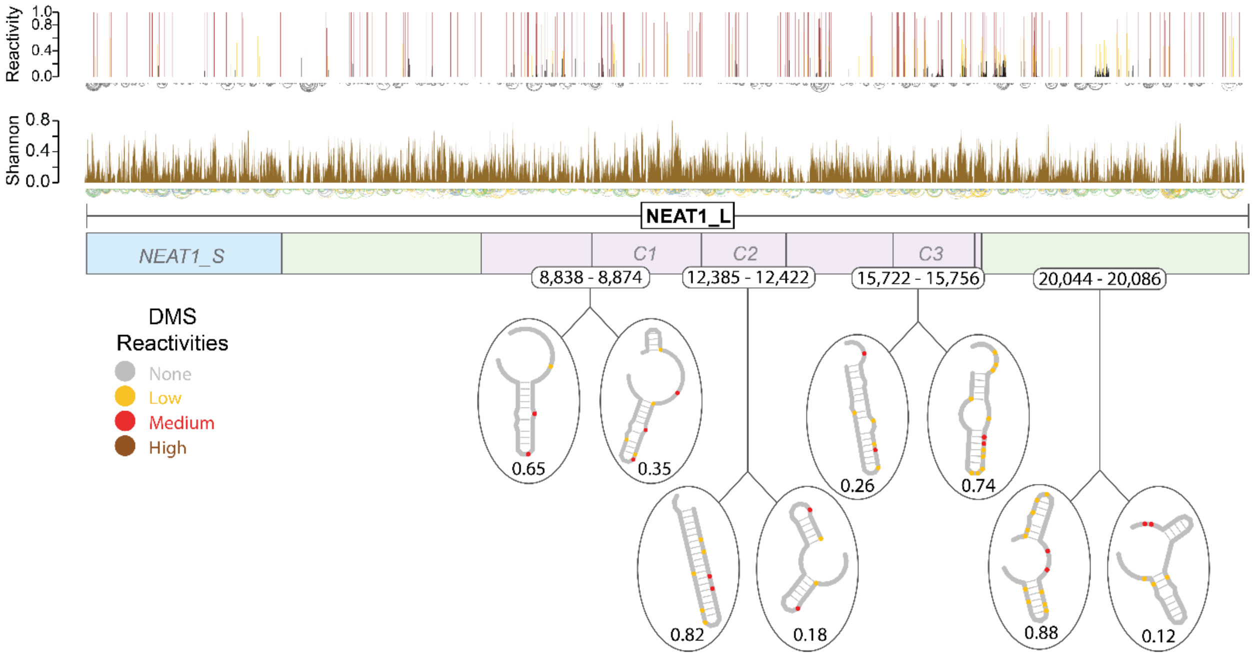


**Supplementary Figure 6.** Attempted structural deconvolution of lncRNA NEAT1 using in cell SHAPE-MaP data against the entire transcriptome. The short window size of DRACO, stemming from the short read lengths of the chemical probing experiments, led to the spurious identification of small, local structurally heterogeneous regions.
